# Inhibition of jasmonate-mediated plant defences by the fungal metabolite higginsianin B

**DOI:** 10.1101/651562

**Authors:** Jean-Félix Dallery, Marlene Zimmer, Vivek Halder, Mohamed Suliman, Sandrine Pigné, Géraldine Le Goff, Despoina D. Gianniou, Ioannis P. Trougakos, Jamal Ouazzani, Debora Gasperini, Richard J. O’Connell

**Affiliations:** UMR BIOGER, INRA, AgroParisTech, Université Paris-Saclay, Thiverval-Grignon, France; Department of Molecular Signal Processing, Leibniz Institute of Plant Biochemistry, Halle (Saale), Germany; Chemical Biology Laboratory, Max Planck Institute for Plant Breeding Research, Cologne, Germany; Centre National de la Recherche Scientifique, Institut de Chimie des Substances Naturelles ICSN, Gif-sur-Yvette, France; Department of Cell Biology and Biophysics, Faculty of Biology, National and Kapodistrian University of Athens, Greece

**Keywords:** *Colletotrichum*, fungal natural product, higginsianin, jasmonate signalling, JAZ protein, plant chemical biology, plant immunity, proteasome, secondary metabolite

## Abstract

Infection of *Arabidopsis thaliana* by the ascomycete fungus *Colletotrichum higginsianum* is characterised by an early symptomless biotrophic phase followed by a destructive necrotrophic phase. The fungal genome contains 77 secondary metabolism-related biosynthetic gene clusters (BGCs), and their expression during the infection process is tightly regulated. Deleting *CclA*, a chromatin regulator involved in repression of some BGCs through H3K4 trimethylation, allowed overproduction of 3 families of terpenoids and isolation of 12 different molecules. These natural products were tested in combination with methyl jasmonate (MeJA), an elicitor of jasmonate responses, for their capacity to alter defence gene induction in *Arabidopsis*. Higginsianin B inhibited MeJA-triggered expression of the defence reporter *VSP1p:GUS*, suggesting it may block bioactive JA-Ile synthesis or signalling *in planta*. Using the JA-Ile sensor Jas9-VENUS, we found that higginsianin B, but not three other structurally-related molecules, suppressed JA-Ile signalling by preventing degradation of JAZ proteins, the repressors of JA responses. Higginsianin B likely blocks the 26S proteasome-dependent degradation of JAZ proteins because it inhibited chymotrypsin- and caspase-like protease activities. The inhibition of target degradation by higginsianin B also extended to auxin signalling, as higginsianin B treatment reduced IAA-dependent expression of *DR5p:GUS*. Overall, our data indicate that specific fungal secondary metabolites can act similarly to protein effectors to subvert plant immune and developmental responses.

**Highlight:** A diterpene secondary metabolite produced by a fungal pathogen suppresses plant jasmonate defense signalling by preventing the proteasomal degradation of JAZ repressor proteins.

## Introduction

The perception of microbial plant aggressors is mediated by the recognition of pathogen-associated molecular patterns (PAMPs) by plant cell surface receptors, which in turn activates a cascade of PAMP-triggered immune (PTI) responses (Dodds and Rathjen 2010; Zipfel and Robatzek 2010). Downstream of PTI activation, these immune responses are regulated by an interconnected network of phytohormone signalling pathways in which jasmonic acid (JA), ethylene (ET) and salicylic acid (SA) play a central role (Pieterse *et al.*, 2012). Antagonistic and synergistic interactions between these pathways provide an additional layer of regulation in which hormone cross-talk allows the plant to fine-tune its immune responses to particular pathogens (Bigeard *et al.*, 2015, Pieterse *et al.*, 2012). A broad range of microbes target these hormones signalling pathways using secreted protein or small molecule effectors in order to manipulate or circumvent plant immunity (Plett et al. 2014; Patkar *et al.*, 2015; Gimenez-Ibanez et al. 2016; Katsir et al. 2008; Groll et al. 2008; Stringlis *et al.*, 2018).

The ascomycete fungus *Colletotrichum higginsianum* causes anthracnose disease in numerous wild and cultivated members of the Brassicaceae, including *Arabidopsis thaliana*. The latter interaction provides a model pathosystem in which both partners are amenable to genetic manipulation and rich genetic resources are available for the plant host. Infection of *A. thaliana* by *C. higginsianum* is characterised by an early symptomless biotrophic phase followed by a destructive necrotrophic phase (O’Connell *et al.*, 2004). As with other hemibiotrophic pathogens, it is assumed that during the biotrophic phase the fungus manipulates living host cells to evade plant defences, while fungal toxins and degradative enzymes are secreted in the necrotrophic phase to kill host cells and mobilise nutrients (Collemare *et al.*, 2019). We previously reported that *C. higginsianum* tightly regulates the expression of secondary metabolism biosynthetic gene clusters (BGCs) at different stages of the infection process (Dallery *et al.*, 2017). Remarkably, no fewer than 14 BGCs are specifically induced early, during penetration and biotrophic colonization, whereas only five are preferentially activated during necrotrophy. Hence, not including possible biosynthetic intermediates, up to 14 different secondary metabolites are potentially delivered to the first infected host cell, where they may contribute to establishing a biotrophic interaction with *A. thaliana*. The transient production of these fungal metabolites exclusively *in planta* presents a major challenge to their structural characterization and functional analysis. In the past decade, deleting proteins involved in shaping the chromatin landscape has allowed the isolation of numerous novel metabolites from diverse axenically grown fungi (e.g. Bok *et al.*, 2009, Fan *et al.*, 2017, Studt *et al.*, 2016, Wu *et al.*, 2016). Recently, we reported a Δ*cclA* mutant of *C. higginsianum* affected in the trimethylation of histone proteins at H3K4 residues which overproduces 12 different metabolites belonging to three terpenoid families, including five new molecules (Dallery *et al.*, 2019a, Dallery *et al.*, 2019b).

Despite the huge efforts made in recent years to characterise the natural products produced by plant-associated microorganisms, to date most studies have only reported on their antimicrobial activity or phytotoxicity and have neglected their potential activity against components of PTI and hormone signalling (Collemare *et al.*, 2019). Indeed, only 30 chemical screens relating to plant biology have been reported in the literature, of which nine tested activity on plant immunity and only one concerned JA signalling (Meesters *et al.*, 2014, Serrano *et al.*, 2015). Using a forward chemical genetic screen, we here identify a fungal natural product that suppresses JA-mediated plant defences. Using different JA-reporter lines in *Arabidopsis*, we show that higginsianin B, a terpenoid metabolite produced by *C. higginsianum*, can prevent the MeJA-dependent degradation of JAZ repressor proteins. Three structural analogues of higginsianin B were found to lack this activity, providing clues to the structure-activity relationship and suggesting candidate functional groups which could help in identifying target binding sites. We also found that the active metabolite is able to inhibit the plant developmental signalling pathway mediated by auxin. Finally, we present evidence that higginsianin B is likely to exert its activity through inhibition of the 26S proteasome. Taken together, our work highlights the importance of fungal secondary metabolites in manipulating plant hormone signalling.

## Methods

### Biological materials

The *Colletotrichum higginsianum* wild-type (WT) strain (IMI 349063A) was maintained on Mathur’s medium as previously described (O’Connell *et al.*, 2004). *Arabidopsis thaliana* accession Columbia (Col-0) was used as the WT line and served as genetic background for the previously described reporters used in this study: *VSP1p:GUS* (Zheng *et al.*, 2006), *PR1p:GUS* (Shapiro and Zhang 2001), *CaMV35Sp:JAZ1-GUS* (Thines *et al.*, 2007), *CaMV35Sp:Jas9-VENUS-NLS* (Larrieu *et al.*, 2015), *JAZ10p:GUSPlus* (Acosta *et al.*, 2013), and *DR5p:GUS* (Ulmasov *et al.*, 1997). Unless otherwise specified, *Arabidopsis* was grown axenically in half-strength Murashige and Skoog (MS) medium (0.5 × MS, 0.5 g∙L^−1^ MES hydrate, pH 5.7). For solid medium, agar was added at 0.7% and 0.85% for horizontal and vertical growth, respectively.

### Chemicals

*C. higginsianum* compound fractions were generated by purifying crude culture extracts using flash chromatography. The pure secondary metabolites used in this study, namely the diterpenoids higginsianin A, B, C and 13-*epi*-higginsianin C, were isolated and structurally identified as previously reported (Dallery *et al.*, 2019b). All fractions and pure compounds were dissolved in dimethyl sulfoxide (DMSO) as stock solutions.

### Quantitative assay for inhibition of JA and SA responses

Hydroponically grown 12-day-old transgenic *Arabidopsis* seedlings of *VSP1p:GUS* and *PR1p:GUS* reporters were used to identify compounds interfering with jasmonate-, or salicylic acid-mediated defences, respectively. Seedlings were treated with compounds for 1 h before inducing reporter gene expression with MeJA (100 µM) or SA (200 µM) dissolved in DMSO. After 24 h, the liquid medium was removed carefully from the wells with a vacuum pump. Seedlings were incubated with 150 µL lysis buffer containing 50 mM sodium phosphate, pH 7.0, 10 mM EDTA, 0.1 % Triton X-100 and 1 mM 4-methylumbelliferyl-β-D-glucuronide (4-MUG; 69602, Sigma-Aldrich) at 37°C for 90 min. The reaction was stopped by adding 50 µL of 1 M Na_2_CO_3_ and 4-MU fluorescence was measured in a microplate reader (excitation/emission wavelength 365/455 nm). Activity was expressed as relative light units (RLU). Each treatment was performed on 5 independent seedlings.

### Histochemical GUS staining

Samples were fixed in 90 % acetone on ice for 1 h, washed in 50 mM NaPO_4_ buffer pH 7.0, vacuum-infiltrated with GUS substrate solution [50 mM NaPO_4_ buffer, pH 7.0, 0.1 % (v/v) Triton X-100, 3 mM K_3_Fe(CN)_6_, 1mM 5-bromo-4-chloro-3-indolyl ß-D-glucuronide] and incubated at 37°C for 2h. Staining was stopped with 70 % ethanol and samples were mounted in 70 % glycerol for observation with a binocular microscope.

### *In vivo* Jas9-VENUS degradation

Inhibition of JAZ protein degradation upon MeJA treatment was assayed using the *Arabidopsis* JA-Ile sensor *CaMV35Sp:Jas9-VENUS-NLS* (Larrieu *et al.*, 2015). After seed stratification for 2 days at 4°C, seedlings were grown vertically for 5 days. Growth conditions were 21°C with a photoperiod of 14h light (100 µE∙m^−2^∙s^−1^). Seedlings were pre-treated with either mock (DMSO in 0.5× MS) or the compound under analysis (30 µM) in a sterile dish for 30 min, then samples were mounted in 60 µL of 30 μM MeJA in 0.5 × MS on microscope slides and imaged immediately (0 min) and 30 min after MeJA treatment. In this way, expression of the reporter was evaluated in individual seedling roots (n = 10 for each condition). To ensure that pre-treatments did not cause reporter degradation, a full sample set was also pre-treated directly on microscope slides and imaged at 0 min and after 30 min. VENUS fluorescence in living roots was imaged with a Zeiss LSM 700 confocal laser scanning microscope with 488 nm excitation and 490-555 nm emission wavelength. All images shown within one experiment were taken with identical settings. Image processing was done with FIJI (http://fiji.sc/Fiji).

### Monitoring Jas9-VENUS degradation by immunoblot

Five-day-old seedlings were grown horizontally in axenic conditions on a nylon mesh (200 µm pore size) supported on MS solid medium. Growth conditions were 21°C with a photoperiod of 14h light (100 µE∙m^−2^∙s^−1^). Pre-treatment and treatment of seedlings was performed as described for microscopy, except that treatments were performed in sterile dishes. E-64, a highly selective cysteine protease inhibitor (E3132, Sigma-Aldrich) and epoxomicin, a specific proteasome inhibitor (E3652, Sigma-Aldrich) were used as controls. Seedlings were snap-frozen in liquid nitrogen and kept frozen for disruption using 3 mm diameter tungsten beads in a Qiagen TissueLyser II operating at 30 Hz, 2 × 1 min. Total proteins from 120 seedlings were extracted with 150 µL of extraction buffer (50 mM Tris-HCl pH 7.4, 80 mM NaCl, 0.1 % Tween 20, 10 % glycerol, 10 mM dithiothreitol, 2× Protease inhibitor cocktail [11873580001, Roche], 5 mM PMSF). Prior to protein quantification, debris were removed by centrifugation at 14,000 rpm, 10 min. Total proteins (40 µg) were separated using SDS-PAGE (10 % acrylamide) and then blotted onto nitrocellulose membranes (1620112, Biorad). Jas9-VENUS and ACTIN were detected using mouse monoclonal antibodies anti-GFP 1:1,000 (11814460001, Roche) or anti-actin 1:2,000 (A0480, Sigma-Aldrich), respectively. The secondary antibody was an anti-mouse coupled to HRP 1:10,000 (W4021, Promega). Detection was performed with the Pico Plus system (34580, Thermo Scientific) and X-ray films (47410 19284, Fujufilm).

### Wounding assays

Horizontally-grown 5-day-old *JAZ10p:GUSPlus* reporter seedlings were pre-treated with either 30 µM DMSO (mock) or 30 µM higginsianin B in water 30 min prior to mechanical wounding one cotyledon as described by Acosta *et al.*, (2013). Pre-treatment was performed by applying 0.5 µL of test solutions to both cotyledons of all seedlings. Histochemical GUS staining was performed 2h after wounding (n = 60 per condition). Alternatively, 1 h after mechanical wounding of one cotyledon, the shoots and roots were collected separately for qRT-PCR analysis of *JAZ10* expression as described previously (Acosta *et al.*, 2013). RNA and cDNA were prepared as in Gfeller *et al.*, (2011). Quantitative RT-PCR was performed as described in Chauvin *et al.*, (2013) using the primers for *JAZ10* (At5g13220) and *UBC21* (At5g25760) previously reported in Gfeller *et al.*, (2011).

### *In vitro* proteasome activity assays

To assess the direct inhibition of proteasomal subunits by higginsianin B, human new born foreskin (BJs) normal fibroblast cells were lysed by using a lysis buffer containing 0.2 % Nonidet P-40,5 mM ATP, 10 % glycerol, 20 mM KCl,1 mM EDTA, 1mM dithiothreitol and 20 mM Tris, pH 7.6). Protein concentration was determined prior to treatment with increasing concentrations of higginsianin B or one of two known proteasome inhibitors (bortezomib or epoxomicin). Chymotrypsin-like (LLVY) and caspase-like (LLE) activities were determined by recording the hydrolysis of fluorogenic peptides Suc-Leu-Leu-Val-Tyr-AMC and Z-Leu-Leu-Glu-AMC, respectively (excitation 350 nm; emission 440 nm).

### Cell-based proteasome activity assays

Measurement of proteasome peptidase activities following cell exposure to the compounds was performed as described previously (Sklirou *et al.*, 2015). Briefly, cells were plated in 60 mm dishes, left to adhere overnight and then treated with the test compounds for 24 or 48 h. The cells were then lysed and proteasome activities were assayed as described above.

### Auxin treatment

Five-day-old *DR5p:GUS* auxin reporter seedlings were grown vertically as described above. Pre-treatment with mock (DMSO in 0.5× MS) or higginsianin B solution (30 µM in 0.5× MS) was performed in sterile dishes for 30 min, followed by 2 h treatment with either mock or naphthaleneacetic acid (NAA, 5 μM in 0.5× MS), a synthetic auxin analogue.

### Statistical Analyses

Statistical analyses were conducted using R software (version 3.4.2) and the packages *Rcmdr* (version 2.4-4) and *conover.test* (version 1.1.5), all available from The Comprehensive R Archive Network (CRAN). The statistical significance of compound treatments on *VSP1p:GUS* and *PR1p:GUS* activation was performed using the Kruskal-Wallis test followed by multiple comparisons using the Conover-Iman test with Benjamini-Hochberg adjustment of *P*-values for false discovery rate (FDR). All proteasome activity tests were performed at least in duplicate and data were statistically analysed with an ANOVA single factor test.

## Results

### Chemical genetics screens identify an inhibitor of JA signalling

Chemical genetics screens using transgenic *Arabidopsis* lines expressing suitable reporter genes are powerful tools to detect small molecules interfering with components of plant defence and hormone signalling (Meesters and Kombrink 2014, Serrano *et al.*, 2015). To search for such activities among *C. higginsianum* metabolites, we generated a small library of partially purified fractions (F1 – F4) and one pure molecule, namely higginsianin B, isolated from liquid cultures of the *C. higginsianum* Δ*cclA* mutant (Dallery *et al.*, 2019b). These were then screened for potential inhibitory activity against SA- and JA-induced defence responses using transgenic plants expressing the β-glucuronidase reporter under the SA-responsive *PATHOGENESIS RELATED 1* (*PR1*) promoter or the JA-responsive *VEGETATIVE STORAGE PROTEIN 1* (*VSP1*) promoter, respectively (Shapiro and Zhang 2001, Zheng *et al.*, 2006). Seedlings grown hydroponically in 96-well plates were first treated with fungal metabolites before inducing expression of the reporter genes with SA or MeJA, respectively. The use of 4-MUG as GUS substrate allowed the fluorimetric quantification of reporter gene expression in intact plants (Halder and Kombrink 2015).

None of the tested compounds were able to inhibit or enhance the SA-mediated activation of *PR1p:GUS* (Supplementary Figure S1). Although seedlings pre-treated with fraction F4 and higginsianin B showed a higher activation of *PR1p:GUS* compared to the DMSO control, these differences were not significant (adjusted *P*-value = 0.25, Kruskal-Wallis with Conover-Iman test). In contrast, fractions F3 and F4 both reduced the MeJA-dependent inducibility of *VSP1p:GUS* expression, by 14 % and 66 %, respectively, compared to mock pre-treated controls (Figure 1A). Purification of compounds from these two fractions identified higginsianin B as the only active metabolite at a concentration of 30 µM. In agreement with this result, comparison of HPLC chromatograms of fractions F1-F4 showed that higginsianin B was present only in fractions F3 and F4 (Supplementary Figure S2). Control seedlings that were not treated with MeJA (uninduced) displayed only basal activation of *VSP1p:GUS* (8% of the level in induced seedlings, Figure 1A). Using this assay, we also found that higginsianin B reduced *VSP1p:GUS* activation in a dose-dependent manner between 3 and 100 µM, with maximal inhibition of 56% at 100 µM (Figure 1B). Given the pronounced inhibitory effect of higginsianin B on the JA pathway, we investigated this activity further.

**Figure 1.**
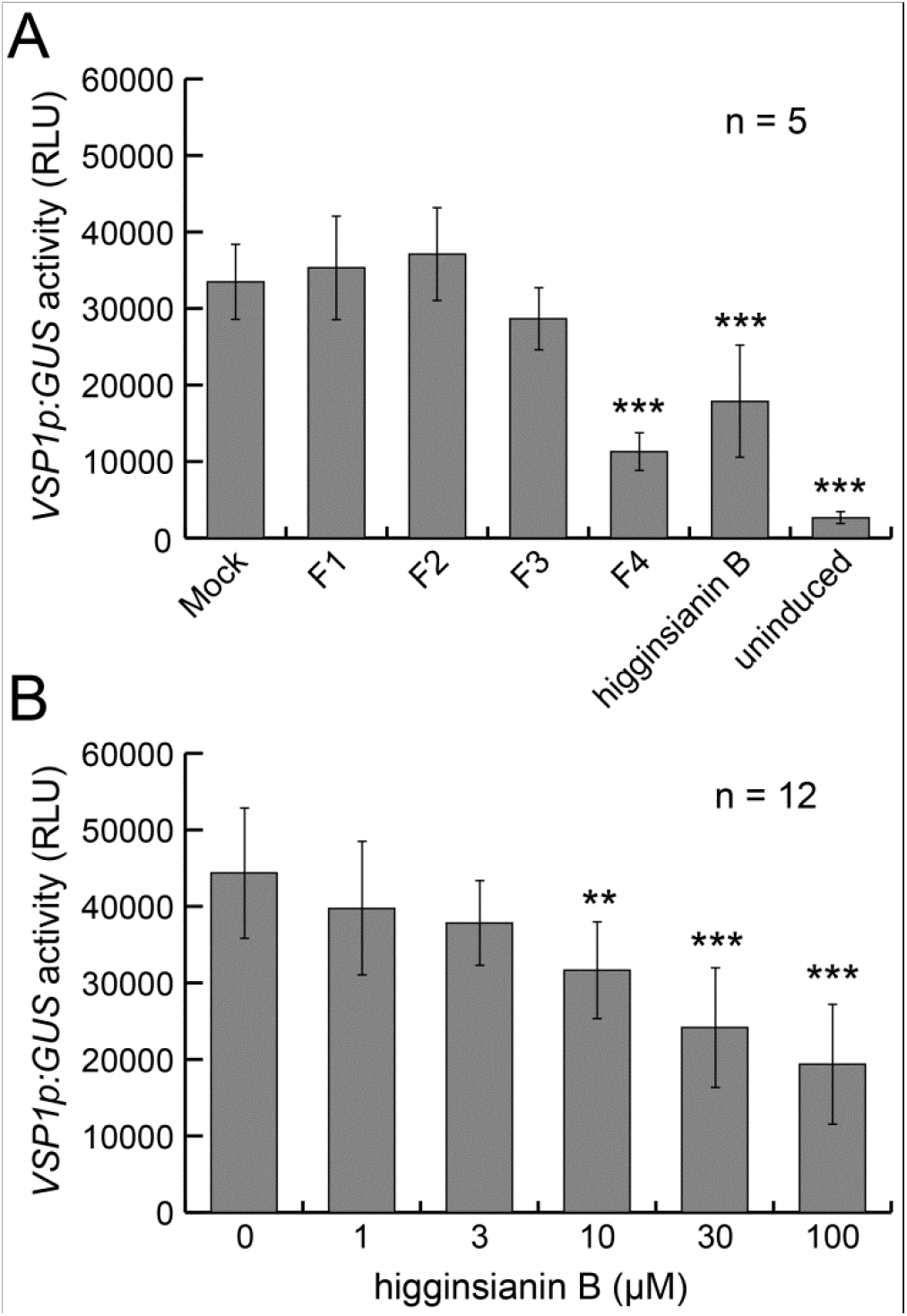
Primary screening identified higginsianin B as a potential inhibitor of JA-mediated plant defence signalling. **(A)** *Arabidopsis* seedlings expressing GUS under the *VSP1* promoter, a marker of JA-mediated plant defences, were pre-treated with metabolite fractions or pure compounds for 1h before MeJA treatment (100 µM for 24 h). Bars represent means *VSP1p:GUS* activity of 5 independent seedlings, ± SD from one representative experiment performed twice. **(B)** Inhibition of *VSP1p:GUS* activity by higginsianin B pre-treatment is dose-dependent. Bars represent means *VSP1p:GUS* activity of 12 independent seedlings, ± SD from one representative experiment performed twice. RLU: Relative Light Unit. **: adjusted *P*-value < 0.01; ***: adjusted *P*-value < 0.001 (Kruskal-Wallis with Conover-Iman test).

### Higginsianin B inhibits JAZ1 degradation

To validate the primary screen result, we tested the effect of higginsianin B on a different marker of the JA pathway, using a transgenic *A. thaliana* line constitutively expressing the JASMONATE ZIM DOMAIN PROTEIN 1 (JAZ1) fused to GUS (*p35S:JAZ1-GUS*) (Thines *et al.*, 2007). JAZ proteins repress JA-responsive genes by binding and inhibiting transcriptional activators such as MYC2 (Pauwels and Goossens 2011). The bioactive jasmonate-isoleucine (JA-Ile) conjugate mediates the binding of JAZ proteins to the F-box protein CORONATINE INSENSITIVE1 (COI1), a member of the Skp1/Cullin1/F-box protein COI1 (SCF^COI1^) complex (Fonseca *et al.*, 2009). JAZ proteins are then poly-ubiquitinated prior to degradation by the 26S proteasome (Chini *et al.*, 2007, Thines *et al.*, 2007). We therefore monitored JAZ1-GUS protein degradation in roots pre-treated with test compounds and then treated with MeJA as described previously (Meesters *et al.*, 2014). While MeJA treatment triggered JAZ1-GUS degradation in mock pre-treated roots, higginsianin B pre-treatment prevented the MeJA-induced degradation of JAZ1-GUS protein at concentrations as low as 0.3 µM and similar to the proteasome inhibitor MG132 (Figure 2) which is known to prevent JAZ1-GUS degradation (Meesters *et al.*, 2014). One possible explanation for this finding is that higginsianin B may inhibit the proteasome-mediated destruction of JAZ1; alternatively, it may block the conversion of inactive MeJA into active JA-Ile. In *Arabidopsis*, this conversion is a two-step process involving a methyljasmonate esterase which produces JA from MeJA and a jasmonoyl-L-amino acid synthetase called JAR1 which converts JA to JA-Ile (Staswick and Tiryaki 2004). When the active JA-Ile was used as inducer in place of MeJA, higginsianin B was still able to inhibit JAZ1-GUS degradation, suggesting that the molecule acts downstream of JA-Ile biosynthesis (Figure 2).

**Figure 2.**
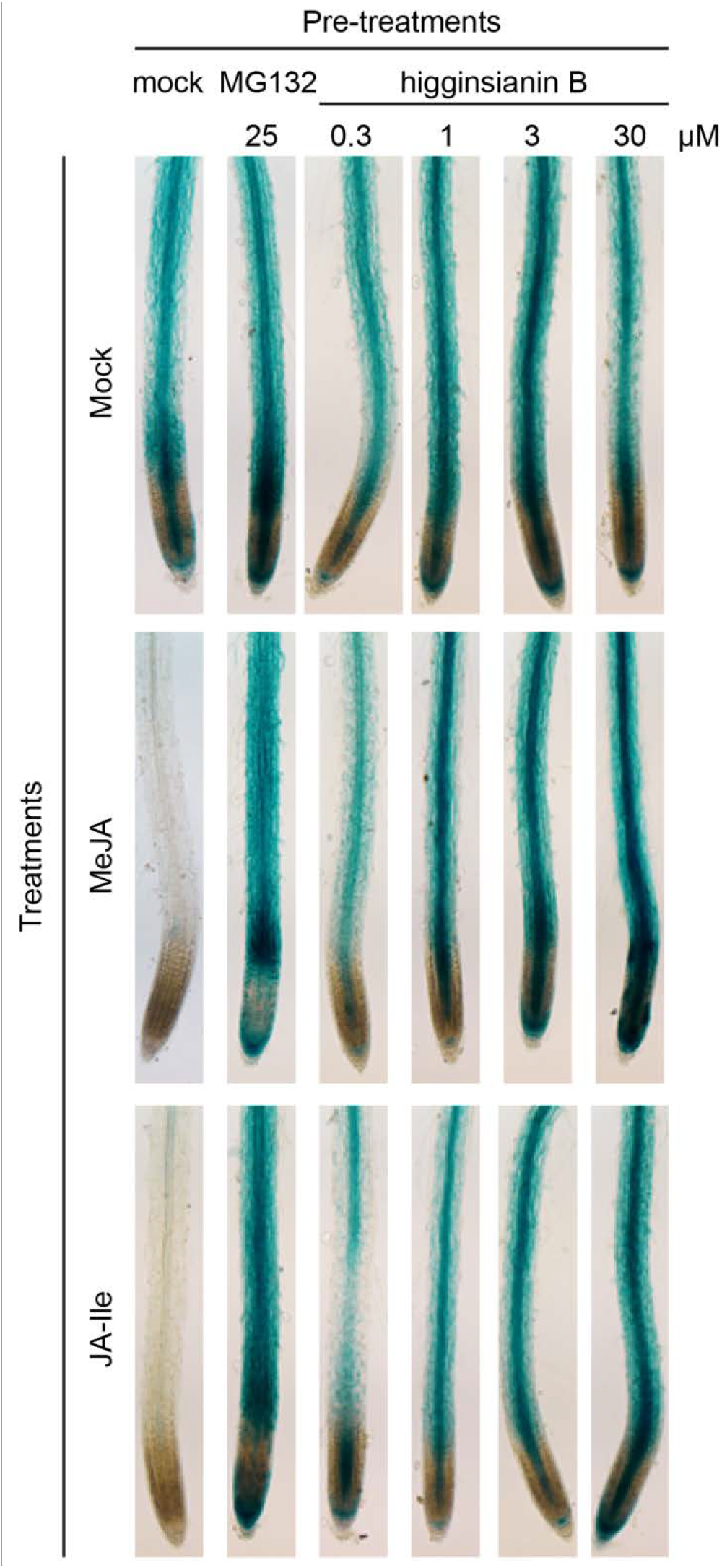
Inhibition of JA-mediated degradation of the JAZ1-GUS protein by higginsianin B. The constitutively expressed JAZ1-GUS chimeric protein is not degraded by mock pre-treatment (60 min) followed by mock treatment (30 min), as shown in seedling roots (upper row) whereas MeJA treatment triggers JAZ1-GUS degradation in mock pre-treated roots (first column, middle row). Pre-treatments with increasing concentrations of higginsianin B prevent MeJA-mediated degradation of chimeric proteins in a dose dependent manner. Using 10 µM of JA-Ile as an inducer instead of 10 µM of MeJA gives similar results indicating that higginsianin B is not inhibiting the conversion of inactive MeJA into the active JA-Ile (lower row). The proteasome inhibitor MG132 was used as a known inhibitor of JAZ1-GUS degradation. Each treatment was performed on at least 5 seedlings and one representative image is presented for each treatment.

### Inhibition of JAZ degradation is specific to higginsianin B

To verify if higginsianin B could inhibit JAZ protein degradation *in vivo*, we monitored its effect on the roots of reporter seedlings constitutively expressing the JA sensor Jas9-VENUS (J9V) consisting of the JAZ9 degron domain (Jas) fused to the VENUS yellow fluorescent protein and a nuclear localization signal (Larrieu *et al.*, 2015). Seedling roots were pre-treated with either mock or compounds under analysis for 30 min, before being treated with MeJA for another 30 min. As expected, MeJA treatment following mock pre-treatment induced J9V reporter degradation, as indicated by the low fluorescence intensity visible in root cell nuclei following the 30 min treatment. (Figure 3A, first row). In contrast, root pre-treatment with higginsianin B (30 µM) strongly inhibited MeJA-induced J9V degradation (Figure 3A, second row). To assess structure-activity relationships, we also tested three other molecules that are structurally related to higginsianin B, namely higginsianin A, higginsianin C and 13-*epi*-higginsianin C (Dallery *et al.*, 2019b). However, pre-treatment with each of these compounds failed to prevent MeJA-induced J9V degradation (Figure 3A), indicating that the inhibitory effect is specific to higginsianin B. By comparing the structures of these molecules (Figure 3B), the functional groups most likely to be required for inhibitory activity are the hydroxyl and / or the 4-isoheptenyl moieties of the bicyclic core.

**Figure 3.**
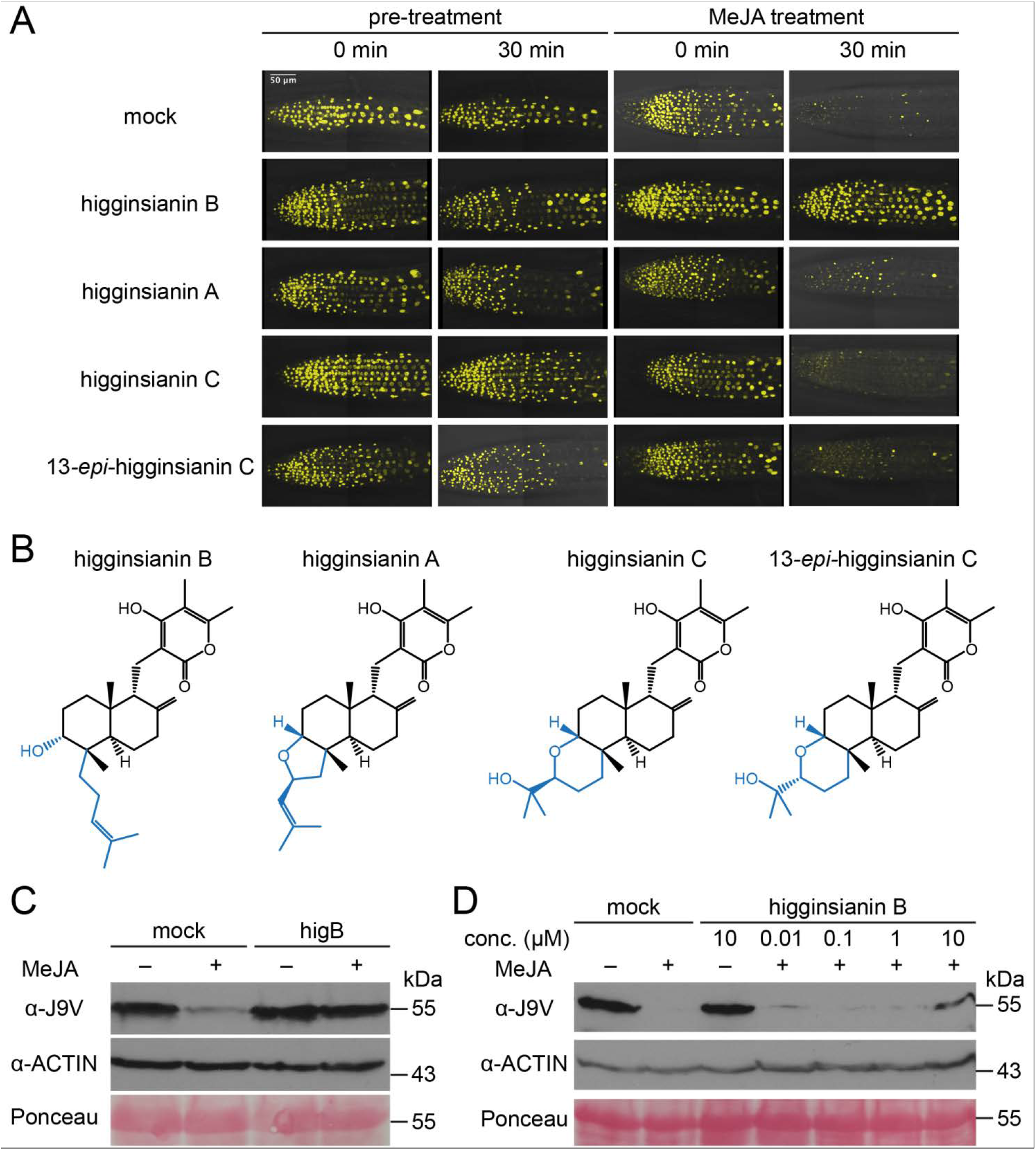
Effect of higginsianin B on Jas9-VENUS (J9V) degradation and structure-activity relationship with other molecules of the higginsianin family. **(A)** Primary roots of the JA sensor J9V before and after pre-treatment with the indicated compounds (30 µM), followed by treatment with MeJA (30 µM). In the control experiment, mock pre-treatment does not induce reporter degradation, while 30 min MeJA treatment is sufficient to induce J9V degradation as indicated by the absence of reporter fluorescence. In contrast, when plants are pre-treated for 30 min with higginsianin B, MeJA treatment is no longer able to promote J9V degradation. Other members of the higginsianin family are unable to prevent MeJA-induced J9V degradation at the tested concentrations (30 µM). **(B)** Chemical structures of higginsianin B, C, A and 13-*epi*-higginsianin C. **(C, D)** Immunoblot analysis of MeJA-induced degradation of J9V (assayed with anti-GFP antibodies). Each lane was loaded with 40 µg of total protein extracts from 60 seedlings. ACTIN (assayed with anti-actin antibodies) and Ponceau S represent loading controls. Protein molecular mass is shown on the right. **(C)** Higginsianin B pre-treatment (30 µM) reduced MeJA-induced Jas9-VENUS degradation. **(D)** Inhibition of MeJA-induced J9V degradation by higginsianin B is dose-dependent.

To further validate results obtained from live-cell imaging, we monitored J9V reporter degradation *in planta* by immunoblot assay. *Arabidopsis* seedlings were pre-treated with either mock or higginsianins for 30 min and subsequently treated with mock or MeJA for 30 min. While MeJA triggered J9V degradation in mock pre-treated seedlings, pre-treatment with higginsianin B at 30 µM prevented J9V degradation (Figure 3C). However, the three other members of this compound family were again inactive at the same concentration (Supplementary Figure S3). A dose-dependency test showed that higginsianin B was active at a concentration of 10 µM (Figure 3D). As controls in this assay, E-64, a highly selective cysteine protease inhibitor was used as an inhibitor of non-proteasomal proteases and epoxomicin as a specific inhibitor of the proteasome. Similar to higginsianin B, epoxomicin inhibited JAS9-VENUS degradation whereas E-64 was inactive (Supplementary Figure S3).

### Higginsianin B inhibits wound-induced JAZ10 activation in roots

So far, our findings revealed that higginsianin B can inhibit JAZ degradation and JA-induced gene expression resulting from exogenous MeJA treatment. To test whether the effect of higginsianin B also extends to suppressing endogenous JA-mediated responses, we assayed JA marker gene expression following mechanical wounding of seedlings pre-treated with higginsianin B. Mechanical wounding of seedling cotyledons is a strong elicitor of JA-dependent gene expression in both shoots and roots, including the activation of the JA-dependent reporter *JAZ10p:GUSPlus* (*JGP*) (Acosta *et al.*, 2013). Pre-treatment of seedling cotyledons with either mock or higginsianin B did not cause reporter activation, while mechanical wounding effectively induced *JGP* expression in wounded shoots in both pre-treatments (Figure 4A). Interestingly, mock pre-treated samples also showed increased *JGP* expression in their roots, whereas higginsianin B pre-treatment reduced the wound-induced reporter activation in this organ (Figure 4A). Quantification of *JAZ10* transcripts further confirmed that higginsianin B pre-treatment reduced wound-induced *JAZ10* accumulation in both shoots and roots as compared to mock treatments (Figure 4B). Furthermore, higginsianin B pre-treatment strongly reduced MeJA-induced *JGP* activation in seedling roots (Supplementary Figure S4A). Taken together, these results indicate that higginsianin B can suppress endogenous JA-mediated responses.

**Figure 4.**
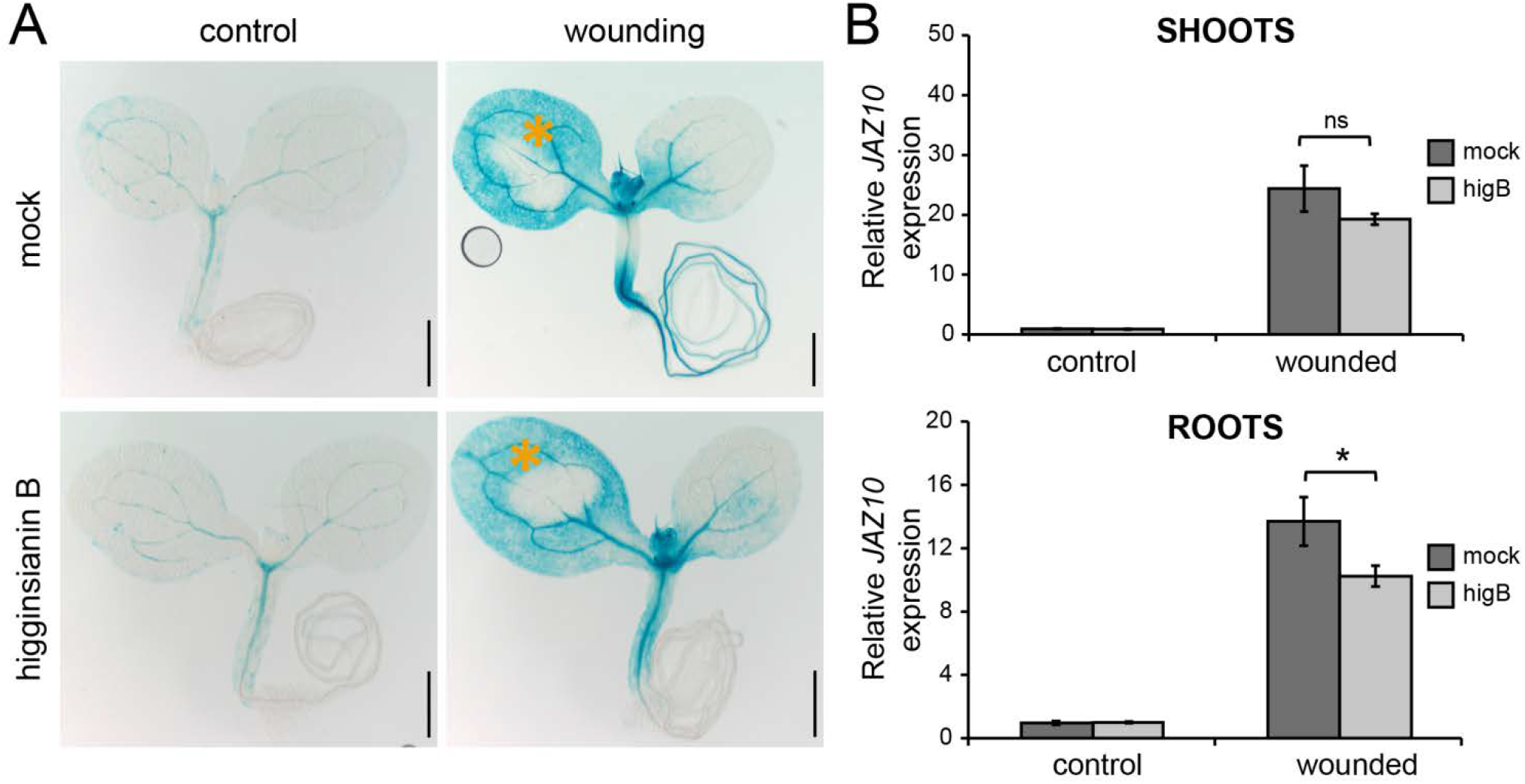
Effect of higginsianin B on wound-induced *JAZ10p:GUSPlus* activation. **(A)** Horizontally grown 5-day old *JAZ10p:GUSPlus* reporter seedlings were pre-treated with mock (30 µM DMSO) or 30 µM higginsianin B by applying 0.5 µL of the pre-treatment solution to their cotyledons for 30 min, after which one cotyledon was mechanically wounded as indicated by orange asterisks. GUS staining was performed 2 h after wounding. Bars = 0.5 mm. **(B)** Quantitative RT-PCR (qRT-PCR) of *JAZ10* expression following 30min pre-treatments with mock or higginsianin B (higB) combined with mechanical wounding. Shoots and roots were collected independently 1 h after wounding aerial organs. *JAZ10* transcript levels were normalised to those of *UBC21* and displayed relative to the expression of mock controls. Bars represent the means of three biological replicates (±SD), each containing a pool of organs from ∼60 seedlings. ns, not significant (*P*-value = 0.08, t-test); *: *P*-value < 0.05 (t-test).

### Higginsianin B affects auxin-mediated signalling

The degradation of JAZ proteins is executed by the 26S proteasome upon poly-ubiquitination by SCF^COI1^ complex (Chini *et al.*, 2007, Thines *et al.*, 2007). Likewise, the 26S proteasome is also involved in auxin perception by co-receptors, the SCF^TIR1/AFB^ ubiquitin ligases and their targets, the AUX/IAA family of auxin response inhibitors (Gray *et al.*, 2001, Tiwari *et al.*, 2001). If higginsianin B blocks JAZ degradation by inhibiting proteasome activity, we reasoned that it may also impact other proteasome-dependent plant responses such as auxin signalling. Treatment of seedling roots with the synthetic auxin naphthaleneacetic acid (NAA) induces expression of the auxin reporter *DR5p:GUS* in the elongation zone (Liu *et al.*, 2017) (Supplementary Figure S4B). Although higginsianin B pre-treatment alone had no any visible effect on the *DR5p:GUS* expression pattern, this pre-treatment not only abolished NAA-mediated reporter induction in the root elongation zone but also reduced *DR5p:GUS* expression in the quiescent centre and root columella (Supplementary Figure S4B). This finding supports the hypothesis that higginsianin B could affect other proteasome-dependent processes, such as the activation of auxin signalling.

### The 26S proteasome is a target of higginsianin B

The impact of higginsianin B on JA- and auxin-mediated signalling pathways suggested the ubiquitin-proteasome system as a possible target. Therefore, to investigate whether higginsianin B can directly inhibit proteolytic activities of the 26S proteasome *in vitro*, human cell lysates containing intact proteasomes were treated with increasing concentrations of the molecule and proteasome activity was measured. Two highly specific proteasome inhibitors, namely bortezomib and epoxomicin, were used as positive controls. We found that higginsianin B inhibited the chymotrypsin-like activity of the proteasome in a dose-dependent manner, with a maximal inhibition of 40% reached at 5 µM; both the bortezomib and epoxomicin were more active in this assay (Figure 5A). Higginsianin B also inhibited the caspase-like proteasomal activity at concentrations of 1 and 5 μΜ, similar to the level of inhibition achieved with epoxomicin and bortezomib (Figure 5B). To measure the effect of higginsianin B on proteasome activities in cell-based assays, we used normal human diploid fibroblasts (BJ cells). In cells treated for 24 h or 48 h with higginsianin B the compound reduced both chymotrypsin-like and caspase-like activities in a dose-dependent manner. The chymotrypsin-like activity was reduced to ~60% at 24 h and ~50% at 48 h relative to the control (Figure 5C). Caspase-like activity was strongly reduced to 35% of the control at 24 h, but only to 70% of the control at 48 h (Figure 5D). Overall, these results suggest that higginsianin B is a potent inhibitor of proteasome proteolytic activities.

**Figure 5.**
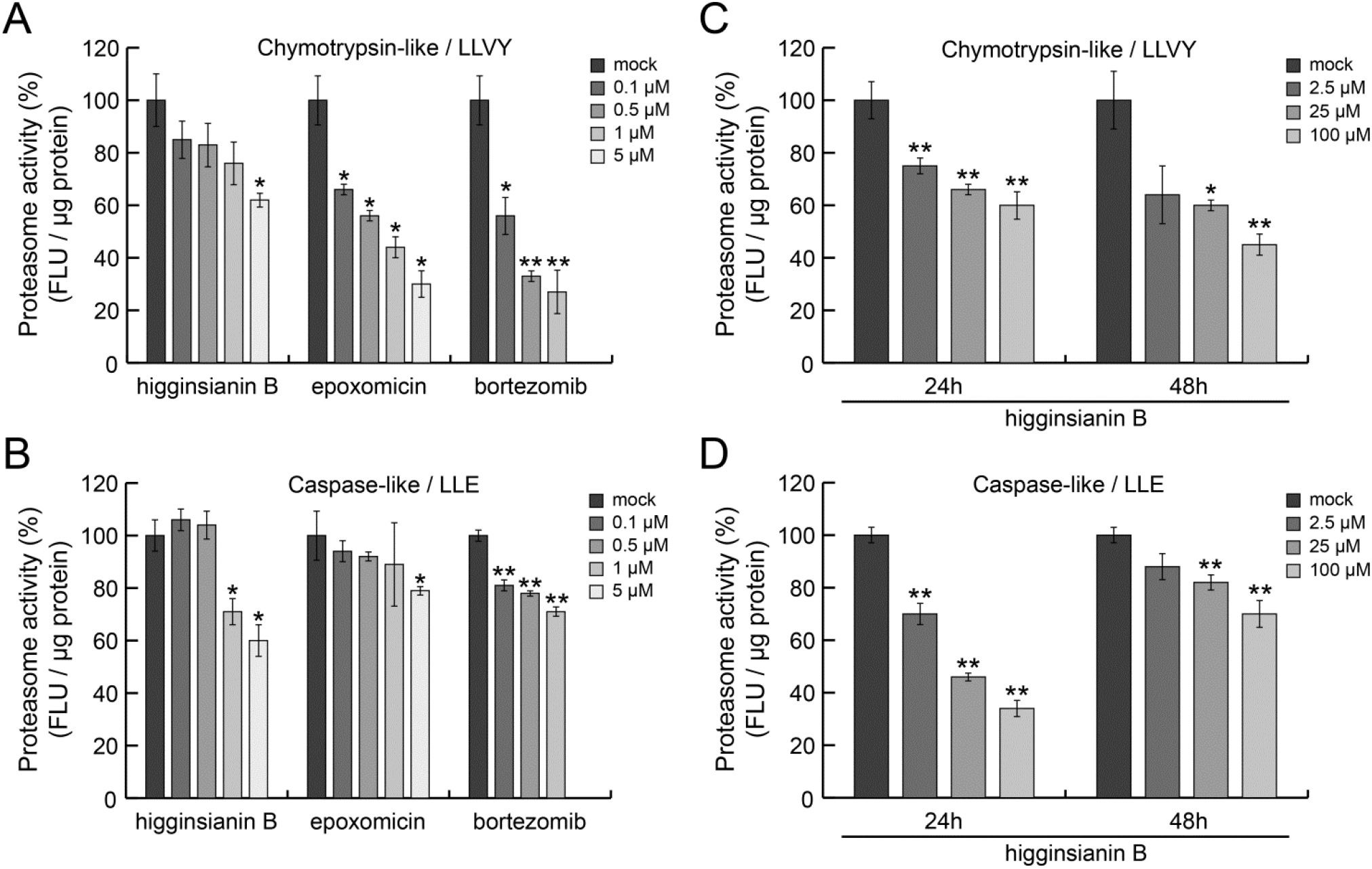
Histograms of inhibition of 26S proteasome activities. **(A, C)** Chymotrypsin-like activity. **(B, D)** Caspase-like activity. **(A, B)** *In vitro* direct inhibition of chymotrypsin-like (panel A) and caspase-like (panel B) activities in a dose-dependent manner by higginsianin B and two known proteasome inhibitors, i.e. epoxomicin and bortezomib. **(C, D)** Cell-based assays showing dose-dependent inhibition of chymotrypsin-like (panel C) and caspase-like (panel D) proteasomal activities in BJ cells exposed for 24 h and 48 h to higginsianin B. Data points correspond to the mean of the independent experiments and error bars denote standard deviation (SD). FLU, Fluorescence unit. *: *P*-value < 0.05; **: *P*-value < 0.01 (ANOVA test).

## Discussion

To date, few chemical genetic screens have been used to systematically search for molecules interfering with components of plant immunity (Dejonghe and Russinova 2017, Serrano *et al.*, 2015). The first small molecule found to inhibit JA-mediated responses in a chemical screen was Jarin-1, a plant-derived alkaloid that was subsequently shown to specifically inhibit the activity of JA-Ile synthetase JAR1, thereby blocking the conversion of JA into bioactive JA-Ile (Meesters *et al.*, 2014). Adopting a similar approach combined with bioassay-guided purification to screen secondary metabolites produced by the *C. higginsianum* Δ*cclA* mutant, we here identified higginsianin B as a novel inhibitor of jasmonate-induced plant defence gene expression. We showed that this diterpenoid can prevent both the wound-induced activation of jasmonate signalling as well as the activation of this pathway by exogenous MeJA. More precisely, we showed higginsianin B acts downstream of the enzymatic conversion of MeJA into JA-Ile by inhibiting the degradation of JAZ proteins, the key repressors of JA signalling in plants. The degradation of JAZ proteins by the ubiquitin-proteasome system (UPS) is essential for de-repressing plant defence genes regulated by JA signalling (Chini *et al.*, 2007, Thines *et al.*, 2007). We present evidence that higginsianin B directly inhibits two catalytic activities of the 26S proteasome, suggesting the molecule most likely blocks the activation of JA-mediated plant defences by inhibiting the proteasomal degradation of JAZ proteins. In agreement with this proposed mode of action, we show higginsianin B also inhibits another proteasome-dependent process, namely the activation of auxin signalling (Gray et al. 2001).

To gain insight into the structural features of higginsianin B that are required for its activity, we tested the three other known members of this compound family, namely higginsianin A, C and 13-*epi*-higginsianin C. Higginsianin B has a bicyclic core substituted by hydroxyl and 4-isoheptenyl groups. In contrast, the three other molecules have a tricyclic core structure with the third ring being a tetrahydrofuran substituted by either an isobutenyl group in the case of higginsianin A or an isopropanol group in the case of higginsianin C and 13-*epi*-higginsianin C. Remarkably, higginsianin B was the only molecule to show activity in JAZ degradation assays suggesting that either the hydroxyl or the 4-isoheptenyl substituents of the bicyclic core (or both) contribute to the activity of higginsianin B. On the other hand, a second hydroxyl group located on the pyrone ring in all higginsianins is unlikely to contribute to this activity, and is therefore a good candidate for tagging higginsianin B with a fluorescent probe for direct visualization of the active metabolite by live-cell imaging. This group could also be exploited for the covalent immobilization of higginsianin B onto a solid support to search for potential protein targets by affinity purification.

While many natural proteasome inhibitors have been discovered from actinobacteria, few were identified from fungi. These include the peptide aldehyde fellutamide B produced by the marine fungus *Penicillum fellutalum* (Hines *et al.*, 2008) and the TMC-95 family of cyclic peptides from the soil saprophyte *Apiospora montagnei* (Momose and Watanabe 2017). Proteasome inhibitors are currently the subject of intense interest as therapeutic agents for the control of cancer and other diseases (Wang et al. 2018; Tsakiri and Trougakos 2015). In this regard it is interesting to note that higginsianin B was recently shown to have antiproliferative activity against glioma, carcinoma and melanoma cell lines (Cimmino *et al.*, 2016). As a novel proteasome inhibitor, higginsianin B therefore merits further investigation as a lead compound for the development of potential therapeutic applications.

Protein turnover by the ubiquitin-proteasome system (UPS) regulates numerous aspects of plant immunity, from pathogen recognition to downstream defence signalling (Marino et al. 2012), and pathogens have evolved protein and chemical effectors to manipulate the UPS to promote plant colonization (Üstün et al. 2016). For example, *Pseudomonas syringae* pv *syringae* secretes the nonribosomal peptide syringolin A which binds covalently to catalytic subunits of the 26S proteasome to inhibit their activity and suppress plant defences (Groll et al. 2008). Two related bacterial Type 3 (T3) secreted effector proteins, XopJ from *Xanthomonas campestris* pv. *vesicatoria* and HopZ4 from *P. syringae* pv *lachrymans*, both attenuate SA-mediated defence by inhibiting proteasome activity through their interaction with RPT6, the ATPase subunit of the 19S regulatory particle of the 26S proteasome (Üstün et al. 2016). Although we have shown here that higginsianin B can directly inhibit two catalytic activities of the mammalian proteasome, further studies are now needed to determine which components of the plant proteasome are the targets of this fungal metabolite and the nature of their interaction.

In the context of JA-mediated defence, the proteasomal degradation of JAZ repressors is targeted by numerous effectors from both pathogenic and mutualistic microbes. For example, the *P. syringae* T3 effectors HopZ1a and HopX1 both activate JA signalling by targeting JAZ proteins for destruction in the proteasome (Jiang et al. 2013; Gimenez-Ibanez et al. 2014). In contrast, the symbiotic ectomycorrhizal fungus *Laccaria bicolor* suppresses JA-mediated defences by secreting the MiSSP7 effector protein, which directly interacts with JAZ proteins to protect them from degradation in the plant proteasome (Plett *et al.*, 2014). The rice blast fungus *Magnaporthe oryzae* weakens JA-mediated plant defence by secreting the inactive hydroxylated JA (12OH-JA) and a monooxygenase enzyme called Abm that hydroxylates JA and depletes levels of endogenous rice JA (Patkar *et al.*, 2015). However, to our knowledge, higginsianin B is the first example of a small molecule produced by any plant-associated fungus that suppresses plant jasmonate signalling by blocking the degradation of JAZ proteins.

In conclusion, our findings raise the possibility that higginsianin B could function during infection as a chemical effector to suppress JA-mediated defences, which are induced at the necrotrophic phase of *C. higginsianum* infection on *Brassica* and *Arabidopsis* (Narusaka et al. 2004; Narusaka et al. 2006). Work is now ongoing to determine at what stage higginsianin B is produced during infection and to genetically test its contribution to fungal virulence and plant defence suppression.

## Supporting information

Dallery et al_Supporting Information

## Supporting Information

Supplementary Figure S1: Screening assay for modulation of salicylic acid signalling pathway using *PR1p:GUS* transgenic line.

Supplementary Figure S2: HPLC-ELSD comparison of four fractions of an active crude extract of *Colletotrichum higginsianum*.

Supplementary Figure S3: Pre-treatments with compounds structurally related to higginsianin B do not influence the MeJA-induced degradation of the JA sensor J9V.

Supplementary Figure S4: Higginsianin B negatively impacts JA- and IAA-triggered gene expression.

## Acknowledgments

The authors are sincerely grateful to Erich Kombrink for valuable discussions. This work was supported by “Chaire d’Excellence” FUNAPP grant (ANR-12-CHEX-0008-01) from the Agence Nationale de la Recherche to R.J.O. and from the Deutsche Forschungsgemeinschaft (grant GA 2419/2-1) to D.G. The Funders had no role in study design, data collection, analysis and interpretation, or writing of the manuscript.

