## Supplementary material for "Inhibition of jasmonate-mediated plant defences by the fungal metabolite higginsianin B": Dallery et al_Supporting Information

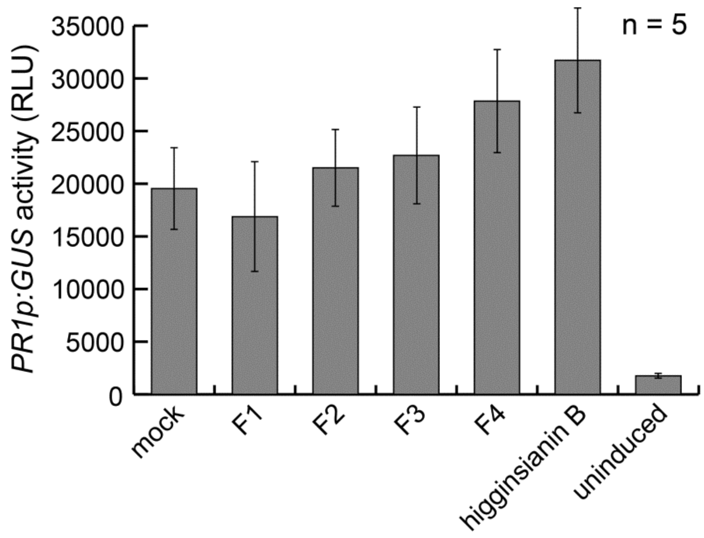


**Supplementary Figure S1 – Screening assay for modulation of salicylic acid signalling pathway.** *Arabidopsis* seedlings expressing GUS reporter under the control of *PR1* promoter, a marker of SA-mediated plant defences, were pre-treated with fractions or pure compound higginsianin B for 1h followed by SA treatment (200 µM) for 24 h. Bars represent means *PR1p:GUS* activity of 5 independent seedlings, ± SD from one representative experiment performed twice. None of the tested fractions or compound were significantly different from the mock control (adjusted P-value = 0.25, Kruskal-Wallis with Conover-Iman test).

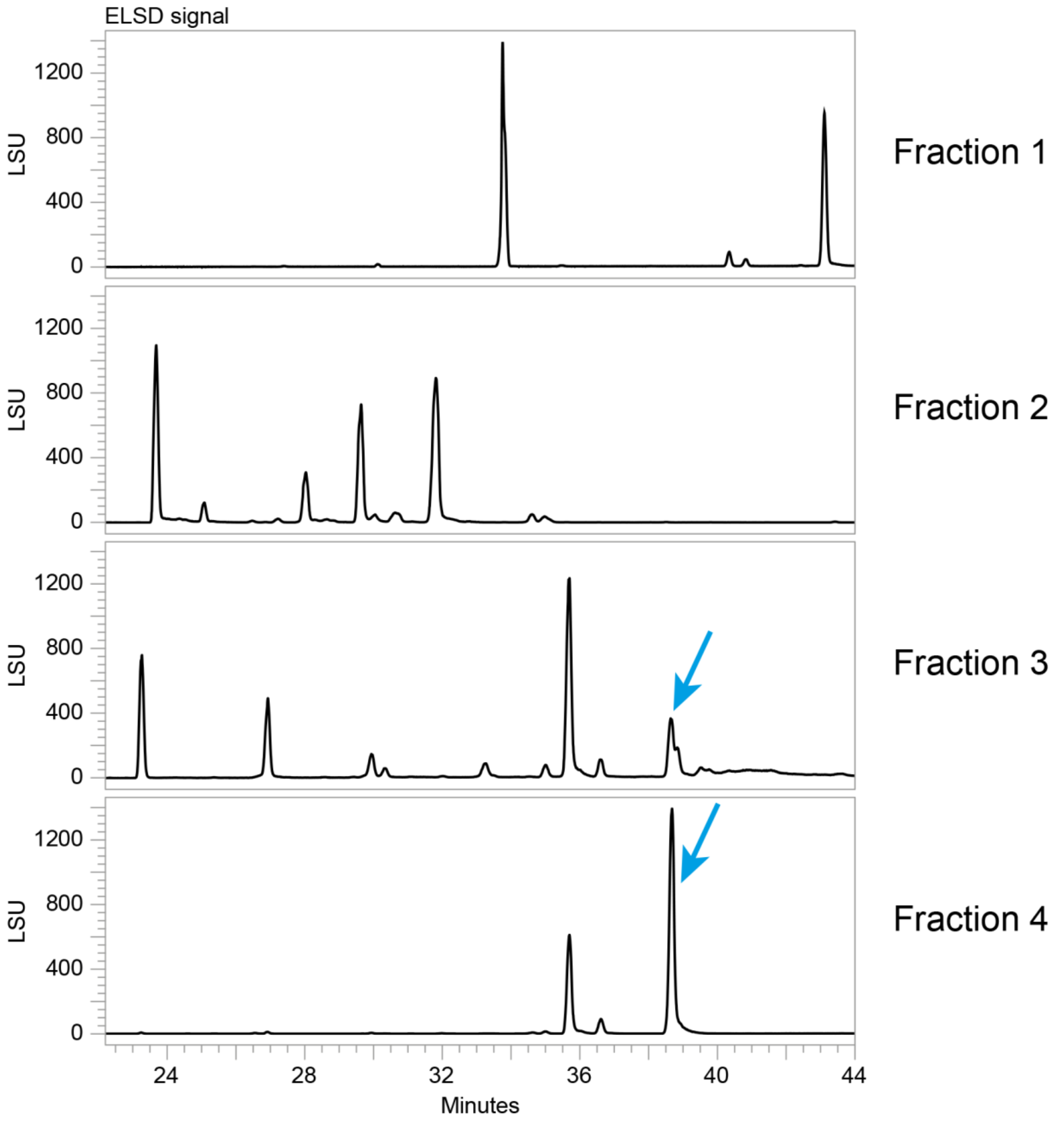


**Supplementary Figure S2 – HPLC-ELSD comparison of four fractions of an active crude extract of *Colletotrichum higginsianum*.** LSU, light scattering unit; ELSD, evaporative light scattering detector. Blue arrows = higginsianin B.

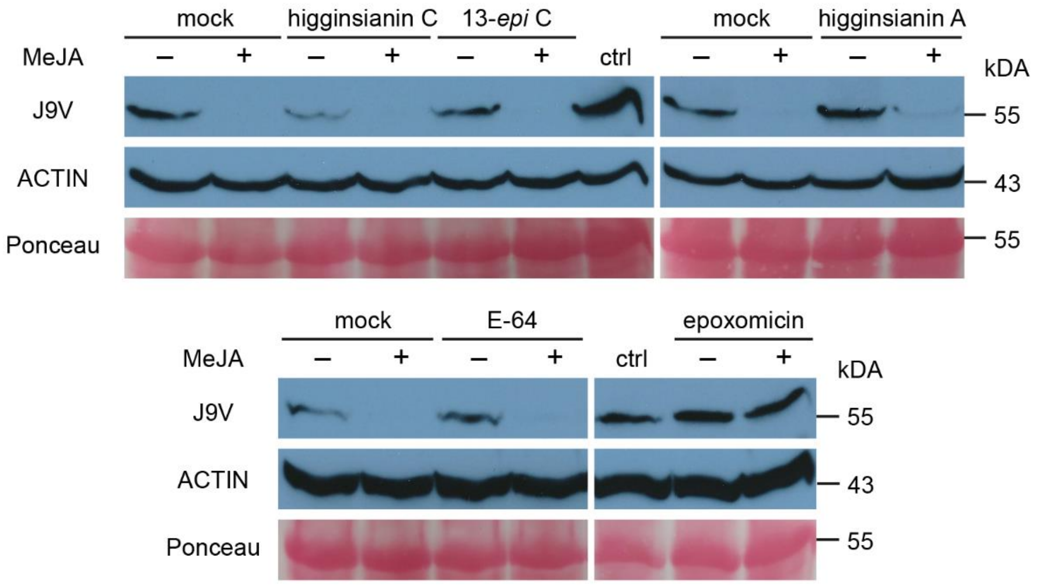


**Supplementary Figure S3 – Pre-treatments with compounds structurally related to higginsianin B (higginsianins A, C and 13-*epi*-higginsianin C) do not influence the MeJA-induced degradation of the JA sensor J9V.** Seedlings were pre-treated with either mock or 30 µM of the indicated compound for 30 min, following with a further 30 min treatment with either mock or 30 µM MeJA. Immunoblots depict 40 µg of total protein extracts from 60 seedlings. J9V was assayed with anti-GFP antibodies; ACTIN (assayed with anti-actin antibodies) and Ponceau S represent loading controls. Protein molecular mass is shown on the right. Higginsianins A, C and 13-*epi*-higginsianin C show very weak activity if any. ctrl refers to untreated Jas9-Venus seedlings. E-64 and epoxomicin, respectively a highly selective cysteine protease inhibitor and a specific proteasome inhibitor, were used as controls. Only epoxomicin could prevent MeJA-induced Jas9-VENUS degradation.

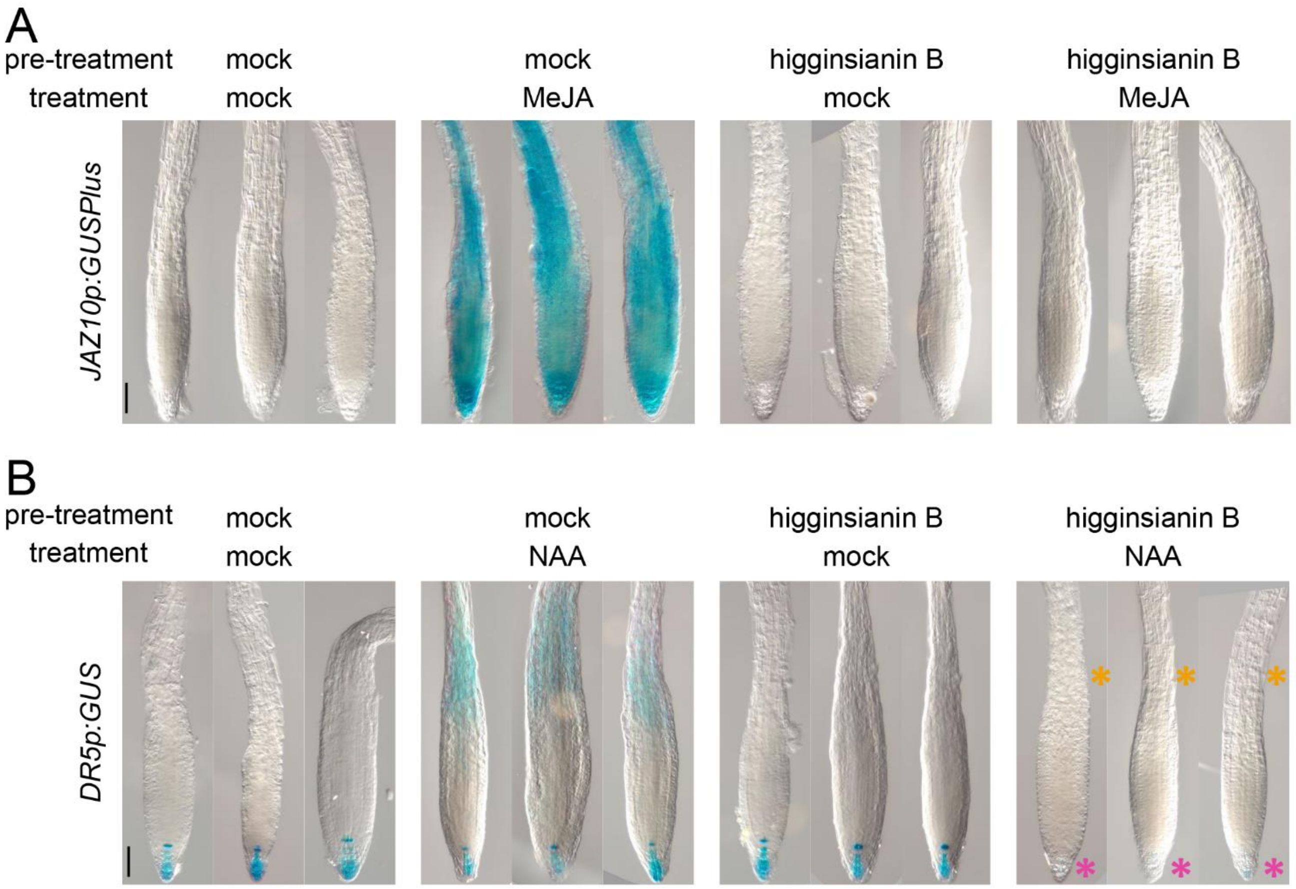


**Supplementary Figure S4 – Higginsianin B negatively impacts JA- and IAA-triggered gene expression.** **(A)** Higginsianin B pre-treatment abolishes the MeJA-mediated induction of *JAZ10p:GUSPlus* in *Arabidopsis* roots. **(B)** Similarly, higginsianin B also inhibits naphthaleneacetic acid (NAA)-mediated induction of the auxin reporter *DR5p:GUS* Note the absence of *DR5p:GUS* staining in the elongation zone of higginsianin B pre-treated / NAA-treated roots (orange asterisks), and reduced reporter expression in the meristem (pink asterisks). Pre-treatments: 30 min (DMSO or 30 µM higginsianin B); Treatments: 2 h. Bars = 50 µm.
